# A Humanized Yeast System for Evaluating the Protein Prenylation of a Wide Range of Human and Viral CaaX Sequences

**DOI:** 10.1101/2023.09.19.558494

**Authors:** Emily R. Hildebrandt, Anushka Sarkar, Rajani Ravishankar, June H. Kim, Walter K. Schmidt

## Abstract

The C-terminal CaaX sequence (cysteine-aliphatic-aliphatic-any of several amino acids) is subject to isoprenylation on the conserved cysteine and is estimated to occur in 1-2% of proteins within yeast and human proteomes. Recently, non-canonical CaaX sequences in addition to shorter and longer length CaX and CaaaX sequences have been identified that can be prenylated. Much of the characterization of prenyltransferases has relied on the yeast system because of its genetic tractability and availability of reporter proteins, such as the **a**-factor mating pheromone, Ras GTPase, and Ydj1 Hsp40 chaperone. To compare the properties of yeast and human prenyltransferases, including the recently expanded target specificity of yeast farnesyltransferase, we have developed yeast strains that express human farnesyltransferase or geranylgeranyltransferase-I in lieu of their yeast counterparts. The humanized yeast strains display robust prenyltransferase activity that functionally replaces yeast prenyltransferase activity in a wide array of tests, including the prenylation of a wide variety of canonical and non-canonical human CaaX sequences, virus encoded CaaX sequences, non-canonical length sequences, and heterologously expressed human proteins HRas and DNAJA2. These results reveal highly overlapping substrate specificity for yeast and human farnesyltransferase, and mostly overlapping substrate specificity for GGTase-I. This yeast system is a valuable tool for further defining the prenylome of humans and other organisms, identifying proteins for which prenylation status has not yet been determined.

**Summary Statement:** We report yeast engineered to express human prenylation enzymes with which prenylation can be investigated for established and novel CaaX sequences associated with proteins involved in human disease.

## Introduction

Post-translational modification of the C-terminal CaaX^1^ sequence is important for regulating the localization and function of many proteins, including the Ras GTPases often cited as archetypical CaaX proteins (Campbell and Philips, 2021; Cox et al., 2015; Ravishankar et al., 2023). CaaX proteins are functionally involved in disease states, such as cancer, Alzheimer’s disease and viral infections, including Hepatitis D and SARS-CoV2 (Jeong et al., 2022; Marakasova et al., 2017; Ring et al., 2022; Soveg et al., 2021; Wickenhagen et al., 2021).

The C-terminal CaaX sequence has been traditionally defined as a Cysteine (C), two aliphatic amino acids (a_1_a_2_), and one of several amino acids (X). The first step in CaaX modification is covalent attachment of a farnesyl (C15) or geranylgeranyl (C20) isoprene lipid to the Cysteine. farnesyltransferase (FTase) generally targets CaaX sequences, while geranylgeranyltransferase-I (GGTase-I) targets the subset of CaaL/I/M sequences (Hartman et al., 2005). CaaX sequences with an aliphatic amino acid at the a_2_ position commonly follow the canonical modification pathway that involves three steps: initial isoprenylation, proteolytic removal of aaX, and carboxymethylation of the exposed prenylated cysteine. In yeast, the Ras2 GTPase and **a**-factor mating pheromone are highly studied examples of canonically modified CaaX proteins. In addition, CaaX sequences lacking a_1_ and a_2_ aliphatic residues can be farnesylated (Kim et al., 2023). These so-called shunted CaaX sequences retain their last three amino acids, resulting in a biochemically distinct C-terminus compared to canonically modified CaaX proteins (Hildebrandt et al., 2016b). The yeast Ydj1 HSP40 protein is a recently characterized example of a shunted CaaX protein.

Farnesylation and geranylgeranylation of CaaX sequences is accomplished by heterodimeric farnesyltransferase (FTase) and geranylgeranyltransferase-I (GGTase-I), respectively. The enzymes share an α subunit while the β subunits provide isoprenoid specificity and substrate recognition (Lane and Beese, 2006; Maurer-Stroh et al., 2003). Human FTase (*Hs*FTase) is composed of *Hs*FNTA and *Hs*FNTB subunits; the orthologous subunits of yeast *Saccharomyces cerevisiae* FTase (*Sc*FTAse) are *Sc*Ram2 and *Sc*Ram1 (**Figure 1A**). Human GGTase-I (*Hs*GGTase-I) is composed of *Hs*FNTA and *Hs*PGGT1B subunits, while the orthologous subunits of yeast GGTase-I (*Sc*GGTase-I) are *Sc*Ram2 and *Sc*Cdc43. In yeast, *RAM1* is not an essential gene (i.e., *ram1Δ* strains are viable), whereas *RAM2* and *CDC43* are essential, raising the possibility that GGTase-I is more critical to yeast life processes relative to FTase.

**Figure 1.**
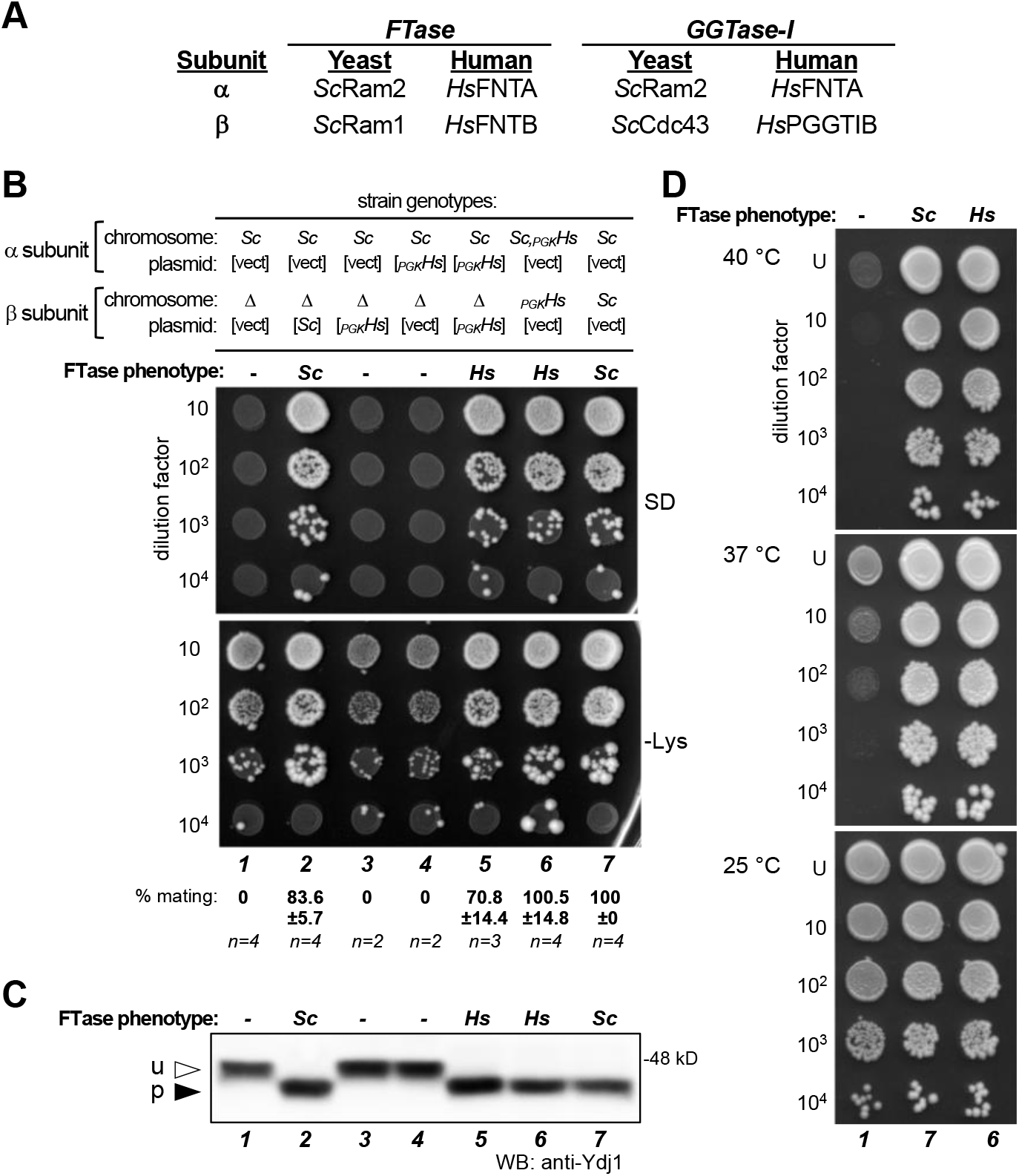
Interspecies complementation studies of FTase subunits. *Hs*FTase and *Sc*FTase subunits are not interchangeable, but co-expression of both α and β subunits of *H*sFTase can restore FTase activity in the absence of *Sc*FTase. **A**) Relationships between yeast and human prenyltransferase subunits. **B**) The mating assay was used to assess FTase dependent production of **a**-factor. *MAT***a** haploid strains were engineered to express yeast (*Sc*) and/or human FTase α and β subunits (*_PGK_Hs*), where subunits were encoded either on plasmids (indicated by brackets) or from chromosomal loci (no brackets). *MAT***a** strains were mixed with *MAT*α cells (SM1068), mixes subject to 10-fold dilutions, and dilution mixtures spotted onto minimal media (SD) and SC-lysine (-Lys) media. Growth on SD indicates diploid formation, which is a direct indicator of **a**-factor mating pheromone production. Growth on -Lys reflects the input of *MAT***a** cells and reflects both unmated haploid and mated diploid cells; unmated haploid cells (*ram1*Δ) grow less well on -Lys due to the absence of FTase activity. Quantitative mating test results are reported below each lane relative to the WT strain (**7**). Strains used were yWS3276 (***1***), yWS3277 (***2***), yWS3278 (***3***), yWS3408 (***4***), yWS3280 (***5***), yWS3282 (***6***), and yWS3283 (***7***). **C**) Gel-shift analysis of Ydj1 using the same strains described in panel **B**. Total cell lysates were prepared and equivalent protein amounts analyzed by Western blot using anti-Ydj1 antibody. u – unprenylated Ydj1; p – prenylated Ydj1. **D**) Thermotolerance test of the FTase-deficient strain compared to wildtype and humanized FTase strains described in panel **B**. Strains used were yWS3276 (***1***), yWS3282 (***6***), and yWS3283 (***7***).

There are 1207 annotated human proteins possessing a CaaX sequence (i.e., Cysteine followed by any 3 amino acids at the C-terminus) within UniProtKB/Swiss-Prot representing 680 unique sequences among the 8000 possible CaaX amino acid combinations. There are also 375 viral proteins possessing a CaaX sequence that could potentially utilize host enzymes for isoprenylation (**Table 1**). Determining the prenylation status of candidate CaaX sequences has depended on a variety of methods and prediction algorithms, each with its own limitations. Methods for case-by-case verification of protein prenylation include radiolabeling with ^3^H-mevalonate, gel mobility analysis, localization studies and mass spectrometry (Anderegg et al., 1988; Hancock et al., 1989; Hildebrandt et al., 2016b; Michaelson et al., 2005; Ravishankar et al., 2023). Methods for systematic verification have relied on metabolic labeling with reagents compatible with click chemistry, peptide arrays, and genetic approaches (Kho et al., 2004; Kim et al., 2023; Rashidian et al., 2013; Storck et al., 2019; Suazo et al., 2018; Suazo et al., 2021; Wang et al., 2014). Differentiating FTase and GGTase-I target specificity has mostly relied on *in vitro* and genetic approaches (Hougland et al., 2010; Kim et al., 2023; Stein et al., 2015). Independent of the methods employed, it remains a distinct challenge to confirm the predictive prenylation of candidate CaaX sequences, especially across species where conservation of target specificity has not necessarily been confirmed.

**Table 1.** Summary of number of proteins identified in UniProtKB/Swiss-Prot database with C-terminal CaaX sequences and with shorter and longer CaaX sequences.

| **Search String** | **Human** | **Non-human Eukaryotes** | **Viruses<sup>a</sup>** | **Total** |
| --- | --- | --- | --- | --- |
| Cxxx> | 1207 | 5047 | 375 | **6629** |
| Cxxxx> (omitted CCxxx>) | 941 | 3582 | 462 | **4985** |
| Cxx> (omitted CCxx>) | 989 | 4206 | 269 | **5464** |
| **TOTAL** | **3137** | **12835** | **1106** | **17078** |

Footnotes for Table 1: <sup>a</sup>Bacteriophage sequences were manually subtracted from the search.

In this study, we have developed yeast strains expressing *Hs*FTase and *Hs*GGTase-I that functionally replace the yeast prenyltransferases in a wide array of tests. Results with a universal reporter for both farnesylation and geranylgeranylation indicate highly overlapping target specificity across species for each prenyltransferase. We also demonstrate the utility of these strains for testing the in vivo prenylation of heterologously expressed human proteins. These humanized strains provide a valuable new resource for protein prenylation research, especially for target specificity studies of the human prenyltransferase in a genetically amenable cell-based system. This resource is expected to enhance the identification and characterization of novel prenyltransferase targets, including those harboring atypical CaaX sequences.

## Results

### Interspecies complementation analysis of FTase subunits

As a key first step toward developing a yeast system to express *Hs*FTase, complementation studies were performed to assess the functional equivalence of yeast and human FTase subunits. Codon optimized genes encoding *Hs*FNTB and *Hs*FNTA were introduced into yeast and evaluated for the ability to complement for loss of yeast FTase β subunit Ram1 (i.e., *ram1Δ)* using an assay that measures production of the farnesylated **a**-factor mating pheromone (**Figure 1B**). No **a**-factor was produced by *ram1Δ* yeast, while complementation with plasmid-encoded Ram1 restored **a**-factor production to near normal levels as measured using an **a**-factor dependent yeast mating assay (i.e., 0% and 83.6% mating relative to WT, respectively). As expected, plasmid encoded *Hs*FNTA did not complement for activity since this strain only carries copies of FTase α subunits (yeast and human). Many human orthologs of yeast proteins can substitute for their yeast counterpart, yet *Hs*FNTB was unable to do so, implying that either the yeast Ram2 and *Hs*FNTB subunits do not interact or a hybrid FTase complex (Ram2-*Hs*FNTB) is non-functional. In fact, **a**-factor was produced only when the human FTase α and β subunits were co-expressed. The observed mating activity was slightly below wildtype levels when *Hs*FTase subunits were plasmid-encoded (70.8% of WT) and at wildtype levels when subunits were integrated into the genome (100.5% of WT). These phenotypes were derived using the strong constitutive phosphoglycerate kinase promoter (i.e., *P_PGK1_*) to drive expression of *Hs*FTase subunits. Expression of *Hs*FNTA and *Hs*FNTB from their orthologous yeast promoters restored mating to less than 1% of wildtype (**Supplemental Figure S1**).

To confirm the ability of yeast expressed *Hs*FTase to fully prenylate other CaaX proteins, the extent of endogenous Ydj1 farnesylation was determined by gel-shift analysis (**Figure 1C**). Farnesylated Ydj1 has faster gel mobility than unfarnesylated Ydj1 when analyzed by SDS-PAGE and immunoblot. Using the same strains evaluated for **a**-factor production, farnesylation of Ydj1 was evident only when *Sc*FTase or *Hs*FTase dimeric complex was present. Farnesylation of Ydj1 by *Hs*FTase was qualitatively complete. When combined with the analysis of **a**-factor production, these results indicate that *Hs*FTase activity is unlikely to be limiting at the cellular level. In addition, we confirmed the ability of chromosomally integrated *Hs*FTase to reverse the temperature sensitive growth phenotype of *ram1*Δ yeast (**Figure 1D**). Together, the results of these complementation studies indicate that *Hs*FTase activity can be engineered to mimic the *Sc*FTase activity levels in a cell-based system.

### *Hs*FTase expressed in yeast modifies human Ras CaaX sequences

Our complementation studies indicated that *Hs*FTase can modify yeast proteins harboring either a canonical CaaX sequence (**a**-factor; CVIA) or a shunted CaaX sequence (Ydj1; CASQ). To extend this observation to other protein contexts and canonical sequences, the farnesylation of the yeast Ras2 GTPase (*Sc*Ras2) was evaluated using the chromosomally integrated *Hs*FTase yeast strain (**Supplemental Figure S2A**). *Sc*Ras2 has 65.6% and 64.4% identity to human NRas and HRas, respectively. In yeast and humans, Ras localization to the plasma membrane depends on canonical modification of the CaaX sequence (Boyartchuk et al., 1997; Michaelson et al., 2005; Ravishankar et al., 2023). Indeed, GFP-*Sc*Ras2 with its natural CaaX sequence (CIIS) was localized to the plasma membrane when *Sc*FTase or *Hs*FTase was expressed (**Figure 2A**; *Sc* and *Hs*, respectively). GFP-*Sc*Ras2 was mislocalized in the absence of FTase activity (*ram1Δ*) or when harboring a CaaX cysteine to serine substitution mutation (SIIS), as has been previously observed (Dong et al., 2003; Ravishankar et al., 2023).

**Figure 2.**
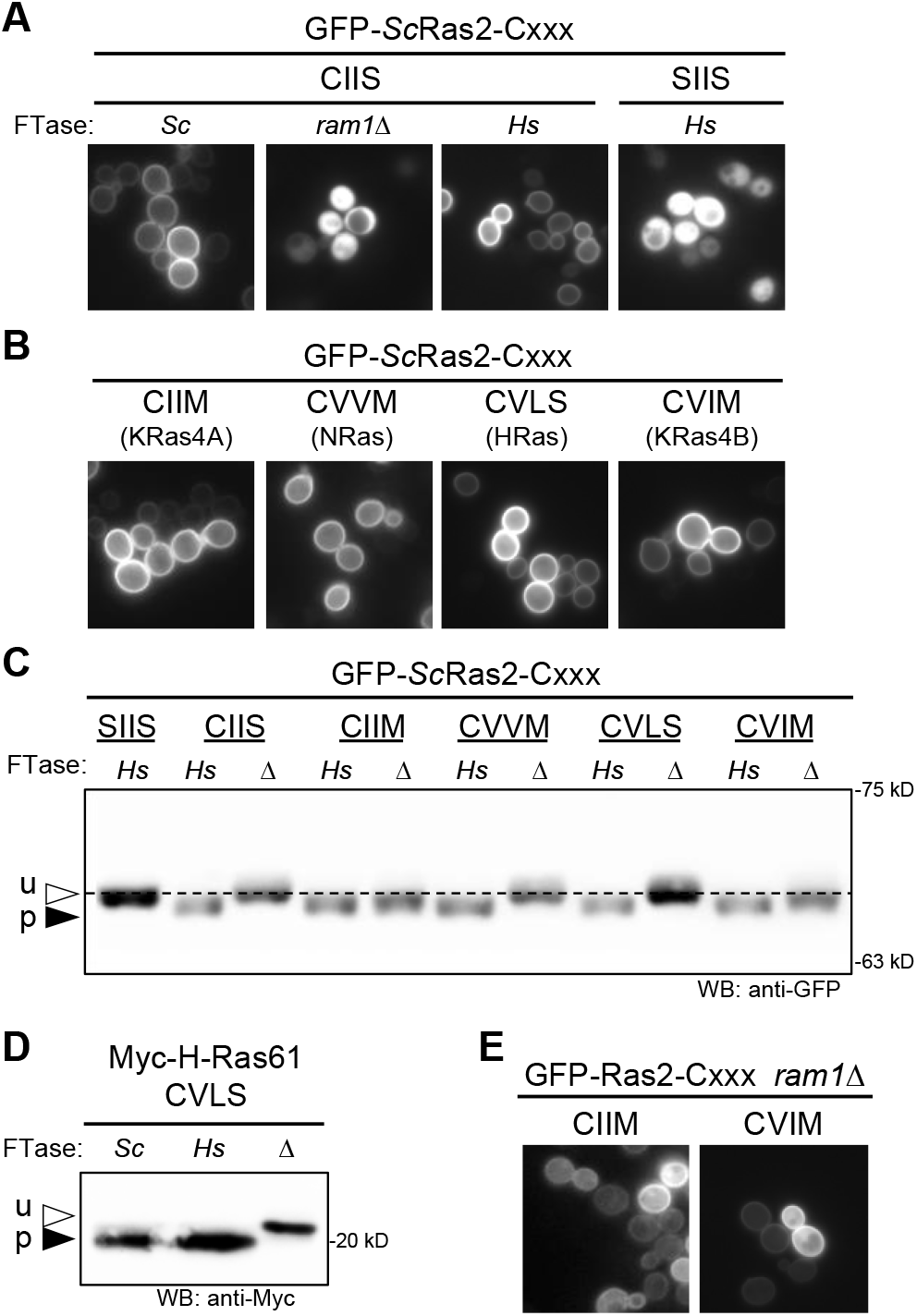
The humanized FTase strain fully modifies yeast and human Ras CaaX sequences. GFP-Ras2 was used as a reporter to evaluate CaaX sequences derived from yeast Ras2 (CIIS) and human proteins KRas4A (CIIM), NRas (CVVM), HRas (CVLS), and KRas4B (CVIM). **A**) The localization of GFP-*Sc*Ras2 produced in wildtype (*Sc*), humanized (*H*s), and FTase-deficient (*ram1Δ*) yeast strains was determined by fluorescence microscopy. GFP-*Sc*Ras2-SIIS is a mutant that cannot be prenylated and is cytosolically localized. **B**) The localization of GFP-*Sc*Ras2-CaaX variants encoding human Ras CaaX sequences produced in the humanized FTase strain was determined by fluorescence microscopy. The source of the CaaX sequence is indicated below each specific sequence. **C**) Western blot analysis of GFP-*Sc*Ras2-CaaX variants produced in humanized (*Hs*) or FTase-deficient (Δ) yeast strains. Total cell lysates were prepared, and equivalent protein amounts analyzed by SDS-PAGE and Western blot using anti-GFP antibody. u –unprenylated GFP-*Sc*Ras2; p – prenylated GFP-*Sc*Ras2. The dashed line was aligned with unprenylated GFP-Ras2 to serve as a visual reference. The plasmids used in panels **A-C** were pWS1735 (GFP-*Sc*Ras2), pWS1889 (GFP-*Sc*Ras2-SIIS), pWS1997 (GFP-*Sc*Ras2-CIIM), pWS1998 (GFP-*Sc*Ras2-CVVM), pWS1999 (GFP-*Sc*Ras2-CVLS), and pWS2000 (GFP-*Sc*Ras2-CVIM). pWS1889 encodes a double cysteine to serine mutation; the upstream cysteine is typically palmitoylated but was mutated to avoid creation of a non-canonical length CaaaX sequence. Yeast strains used were BY4741 (wildtype *Sc*FTase), yWS3220 (*Hs*FTase), and yWS3202 (*ram1Δ*). **D**) Myc-HRas61 (p-05547) was produced in wildtype (*Sc*; yWS2544), humanized FTase (*Hs*; yWS3186), or FTase-deficient (Δ; yWS3209) yeast strains. HRas61 is the Q61L oncogenic derivative of human HRas (Adari et al., 1988; Farnsworth et al., 1991). **E**) The localization of GFP-*Sc*Ras2-CaaX variants (CIIM and CVIM) that undergo alternate prenylation by *Sc*GGTase-I in the absence of FTase (i.e., *ram1Δ* strain background).

We utilized the GFP-*Sc*Ras2 reporter to further evaluate the ability of *Hs*FTase to recognize CaaX sequences associated with human Ras orthologs: *Hs*KRas4A (CIIM), *Hs*NRas (CVVM), *Hs*HRas (CVLS), and *Hs*KRas4B (CVIM) (**Figure 2B**). All the sequences directed GFP-*Sc*Ras2 to the plasma membrane, indicating that they are farnesylated by *Hs*FTase. As additional confirmation of prenylation by *Hs*FTase, the mobilities of GFP-*Sc*Ras2-CaaX variants were analyzed by gel-shift assay (**Figure 2C**). Like farnesylated Ydj1, prenylated *Sc*Ras2 migrates faster relative to unprenylated protein when analyzed by SDS-PAGE. GFP-*Sc*Ras2 exhibited faster mobility relative to protein produced in the absence of FTase (i.e., *ram1Δ*) in the context of its wildtype sequence (CIIS) and two human Ras CaaX sequences (CVVM and CVLS; CIIM and CVIM are discussed below), indicative of complete prenylation. Gel-shift studies were also used to examine the farnesylation of heterologously expressed human oncogenic protein Myc-HRas61 (CVLS) that was heterologously expressed in the humanized yeast strain (Adari et al., 1988; Farnsworth et al., 1991). It too was completely prenylated (**Figure 2D**).

Some CaaX sequences, including those associated with KRas4A (CIIM), KRas4B (CVIM) and NRas (CVVM), are alternately prenylated by *Hs*GGTase-I in the absence of *Hs*FTase activity, which is often established in systems through the action of FTase inhibitors (Mohammed et al., 2016; Whyte et al., 1997). Consistently, GFP-*Sc*Ras2-CaaX variants harboring CIIM and CVIM sequences exhibited mobility shifts of less magnitude in the gel-shift assay (**Figure 2C**), which we attribute to alternate prenylation by yeast GGTase-I in the absence of FTase (*ram1*Δ). To bolster this conclusion, we determined that GFP-*Sc*Ras2-CaaX variants CIIM and CVIM exhibited plasma membrane localization in the FTase-deficient *ram1Δ* strain, which would be predicted if they were geranylgeranylated (**Figure 2E**). Combined, these observations suggest that *Sc*GGTase-I also has the opportunistic ability to modify certain CaaX sequences in the absence of FTase activity (i.e., *ram1Δ*), resulting in geranylgeranylated products having similar although not identical mobility as the farnesylated products.

### *Hs*FTase expressed in yeast modifies the non-canonical human DNAJA2 CAHQ sequence

The ability of *Sc*FTase to modify the non-canonical CaaX sequence associated with Ydj1 (CASQ) and other non-canonical sequences has been reported (Berger et al., 2018; Hildebrandt et al., 2016b; Kim et al., 2023; Ravishankar et al., 2023). The complementation studies described above indicate that *Hs*FTase can also modify the Ydj1 CASQ sequence (see **Figure 1**). To further investigate the breadth of sequences recognized by *Hs*FTase, we used thermotolerance and gel-shift assays to evaluate a set of Ydj1-CaaX variants representing sequences that are well characterized in terms of modification status. In the thermotolerance assay, yeast lacking Ydj1 (vector) or an unmodifiable Ydj1(SASQ) are unable to grow at 40 °C because Ydj1 farnesylation is required for efficient growth at elevated temperatures (**Figure 3A**). All the remaining Ydj1-CaaX variants exhibited growth profiles that were nearly identical whether farnesylation was supported by *Sc*FTase or *Hs*FTase. Yeast expressing Ydj1 with prenylation-only sequences (CASQ and CAHQ) supported the most robust thermotolerance; these sequences were derived from yeast Ydj1 and human DNAJA2, respectively. The prenylation-only status of CAHQ was previously based on homology and prediction algorithms (Hildebrandt et al., 2016b; Kim et al., 2023) and is confirmed here by the observation that CAHQ supports thermotolerance similar to CASQ and is resistant to proteolysis by the CaaX proteases (**Supplemental Figure S3**). Yeast expressing Ydj1 with canonical CaaX sequences (CVIA and CTLM) exhibited partial growth under this condition as has been previously observed (Hildebrandt et al., 2016b); these sequences were derived from yeast **a**-factor and Gγ Ste18, respectively. Side-by-side SDS-PAGE analysis of the Ydj1-CaaX variants revealed that both canonical and prenylation-only CaaX sequences are fully prenylated by both *Sc*FTase and *Hs*FTase (**Figure 3B**). Additional support for the ability of *Hs*FTase to modify these sequences is evident when comparing the mobilities of Ydj1-CaaX variants in the absence of FTase (i.e., *ram1Δ*) (**Figure 3C**).

**Figure 3.**
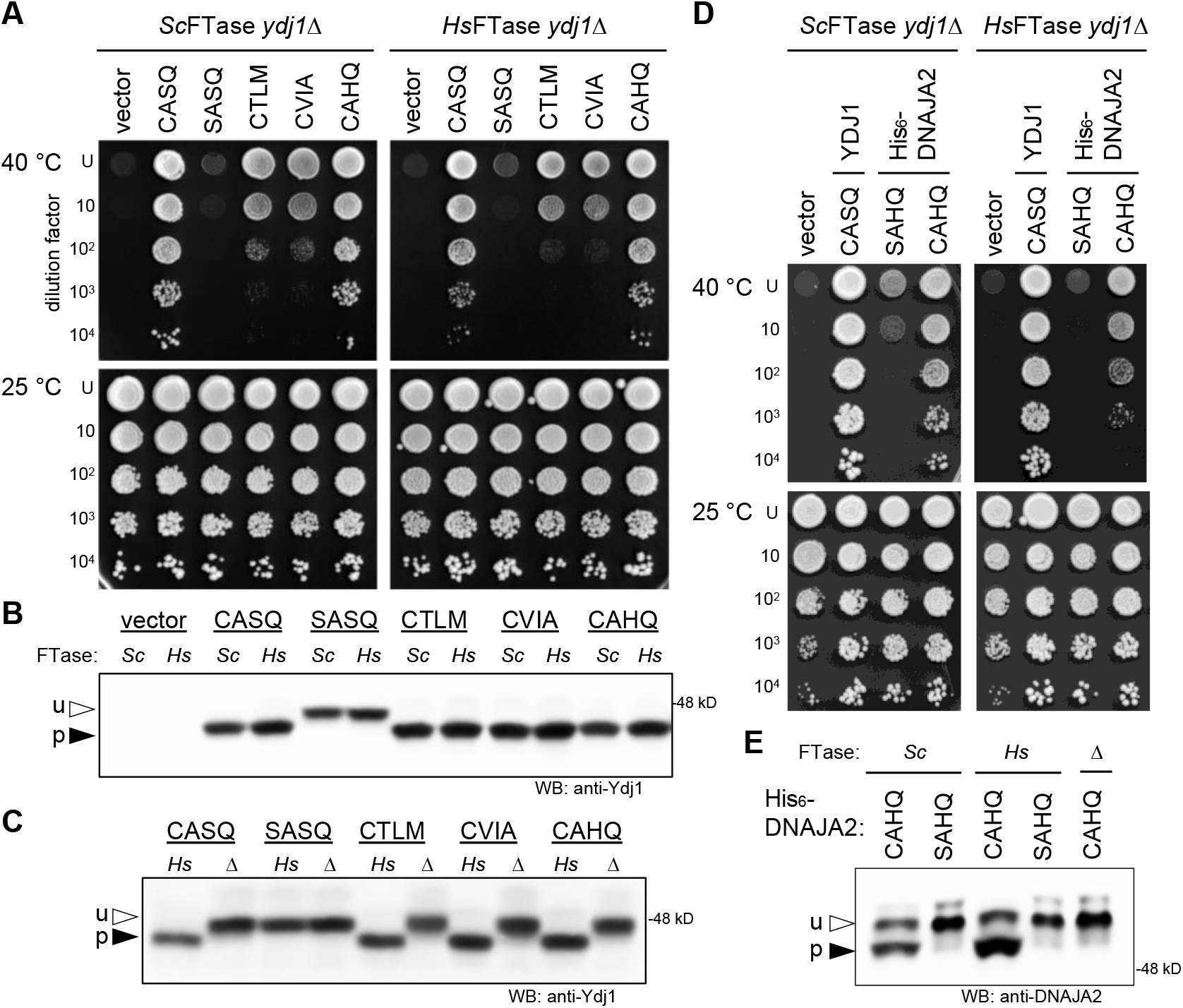
The humanized FTase strain modifies HSP40-CaaX variants. A Ydj1-CaaX reporter was used to evaluate CaaX sequences from yeast HSP40 Ydj1 (CASQ), an unmodifiable sequence (SASQ), yeast Ste18 (CTLM), yeast **a**-factor (CVIA), and human HSP40 DNAJA2 (CAHQ). **A**) The thermotolerance assay was used to determine prenyl modification of Ydj1. Strains were engineered to express *Sc*FTase (left panels) or *Hs*FTase (right panels) and indicated plasmid-encoded Ydj1-CaaX variants. Strains were cultured to equivalent density, cultures subject to 10-fold dilutions, and dilution mixtures spotted onto rich media (YPD) at either room temperature (RT) or elevated temperature (40 °C). Growth at 40 °C reflects farnesylation of Ydj1. **B**) Gel-shift analysis of Ydj1-CaaX variants. Total cell lysates prepared using the strains described in panel **A** were analyzed by SDS-PAGE and Western blot using anti-Ydj1 antibody. u – unprenylated Ydj1; p – prenylated Ydj1. **C**) Comparison of Ydj1-CaaX variant gel mobilities in the absence (Δ) and presence of *Hs*FTase (*Hs*). **D**) Thermotolerance analysis of strains producing native Ydj1 (CASQ), plasmid-encoded human His_6_-DNAJA2 (CAHQ), and an unmodifiable sequence (SAHQ). Strains were evaluated as described in panel **A**. **E**) Gel-shift analysis of human His_6_-DNAJA2 and His_6_-DNAJA2-SAHQ produced in wildtype (*Sc*), humanized FTase (*Hs*), and FTase-deficient (*Δ*) yeast strains. Total cell lysates were prepared and analyzed as described in panel **B**. For all panels the strains used were yWS2544 (*Sc*FTase *ydj1Δ*), yWS3186 (*Hs*FTase *ydj1Δ;* see **Supplemental Figure S2B**), and yWS3209 (*ram1Δ ydj1Δ*). Plasmids used were pRS316 (vector), pWS942 (YDJ1), pWS1132 (YDJ1-SASQ), pWS1246 (YDJ1-CTLM), pWS1286 (YDJ1-CVIA), pWS1495 (YDJ1-CAHQ), pWS1424 (His_6_-DNAJA2), and pWS2256 (His_6_-DNAJA2-SAHQ).

The observations made using Ydj1 as a reporter were also evident for *Hs*DNAJA2 (CAHQ) that was heterologously expressed in the humanized yeast strain. *Hs*DNAJA2 has 46% identity to Ydj1 and can complement some Ydj1-associated defects (Whitmore et al., 2020). *Hs*DNAJA2 supported thermotolerance in the context of both *Sc*FTase and *Hs*FTase strains, and this phenotype was prenylation dependent because DNAJA2-SAHQ did not support thermotolerance (**Figure 3D**). By gel-shift analysis, both *Sc*FTase and *Hs*FTase modified *Hs*DNAJA2 (**Figure 3E**). These results further extend the utility of the humanized yeast strain for studies of heterologously expressed human CaaX proteins. Moreover, these observations support the hypothesis that *Sc*FTase and *Hs*FTase have highly similar sequence recognition profiles that are likely broader than the CaaX consensus sequence.

### *Hs*FTase expressed in yeast modifies a range of human and viral CaaX protein sequences

Studies on the prenylation of CaaX proteins in most systems is hindered by several factors ranging from difficulty in odetection of low abundance proteins, to lack of phenotypic readouts, to labor and time costs for case-by-case studies. Many of these hurdles can be overcome using the Ydj1 reporter and genetically tractable yeast system described in this study. To support the utility of our approach for investigations of *Hs*FTase specificity, Ydj1-CaaX variants harboring the CaaX sequences of known prenylated human proteins were evaluated by gel-shift assay. Each variant was evaluated in the presence and absence of *Hs*FTase to ensure that gel-shifts were due to *Hs*FTase and to assess whether alternate prenylation was possible. The CaaX sequences evaluated were from chaperone proteins (Nap1L1, DNAJA1 and Pex19), chromosome stability proteins (CENPE, CENPF and Spindly), nuclear lamins (Prelamin A and Lamin B), Ras-like proteins (Rab38, KRas4A, KRas4B and HRas) and Prickle1 (**Figure 4A** and **Supplemental Table S5**) (Ashar et al., 2000; Kho et al., 2004; Kim et al., 1990; Leung et al., 2007; Moudgil et al., 2015; Onono et al., 2010; Palsuledesai et al., 2014; Storck et al., 2019; Strutt et al., 2013; Varela et al., 2008). Apart from the Rab38-associated sequence (CAKS), every sequence was completely modified by *Hs*FTase. CAKS is reported to be underprenylated in the context of Rab38, which is consistent with our findings (Kohnke et al., 2013). Sequences ending in methionine (CSIM, CAIM, CLIM, CIIM and CVIM) were fully modified in the presence of *Hs*FTase yet partially modified in the absence of FTase (*ram1Δ*), suggesting that *Sc*GGTase-I contributes to their prenylation. These observations are consistent with studies reporting that *Sc*FTase, *Hs*FTase, and *Hs*GGTase-I recognize CaaM sequences, which we propose is a property that extends to *Sc*GGTase-I as well (Caplin et al., 1994; Zhang et al., 1997).

**Figure 4.**
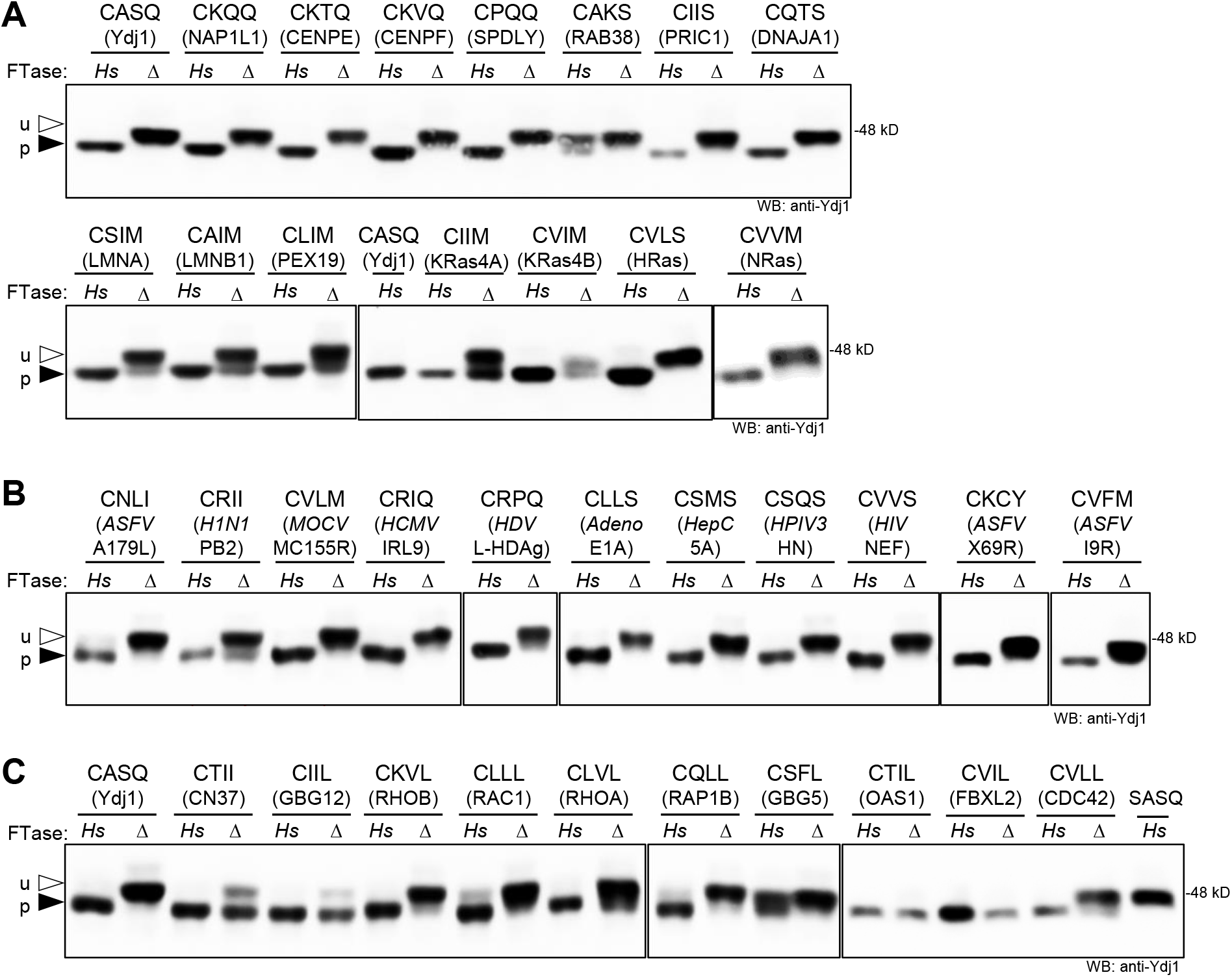
The humanized FTase yeast strain modifies Ydj1-CaaX variants with CaaX sequences occurring in human and viral proteins. Ydj1-CaaX variants were produced in the *Hs*FTase (*Hs*) and FTase deficient (Δ) strains and evaluated by gel-shift assay as described in Figure 3B. The source of the CaaX sequence is indicated below each specific sequence. **A**) Analysis of Ydj1-CaaX variants based on the indicated human proteins. **B**) Analysis of Ydj1-CaaX variants based on mammalian viral proteins. Virus abbreviations are human influenza virus (*H1N1*), Molluscum contagiosum virus (*MOCV*), human cytomegalovirus (*HCMV*), Hepatitis Delta Virus (*HDV*), human adenovirus 1 (*Adeno*), human Hepatitis C virus (*HepC*), human parainfluenza virus 3 (*HPIV3*), Human immunodeficiency virus 1 (*HIV*), and African swine fever virus (*ASFV*). **C**) Analysis of Ydj1 with CaaL/I sequences based on the indicated human proteins. Strains used were yWS3186 (*Hs*) and yWS3209 (Δ). Plasmids used are listed in **Supplemental Table S3**. See **Supplemental Table S1** for gel quantification.

The Ydj1 reporter system was also used to evaluate the prenylation of CaaX sequences associated with human and swine viral proteins (**Figure 4B** and **Supplemental Table S5**). The importance of prenylation in viral infection is an emerging line of inquiry for infectious disease therapy but documentation of viral proteins prenylation is limited (Einav and Glenn, 2003; Marakasova et al., 2017). All of the viral CaaX sequences evaluated were modified by *Hs*FTase. These sequences were associated with influenza virus H1N1 PB2 (CRII); *Molluscum contagiosum* virus subtype 1 MC155R (CVLM); human cytomegalovirus protein IRL9 (CRIQ); Hepatitis Delta Virus L-HDAg (CRPQ); human adenovirus1 E1A (CLLS); hepatitis C virus non-structural 5A protein (CSMS); human parainfluenza virus 3 hemagglutinin-neuraminidase (CSQS); HIV NEF protein (CVVS); and African swine fever virus proteins Bcl-2 homolog A179L (CNLI), X69R (CKCY) and I9R (CVFM). Of these, only hepatitis L-HDAg has been biochemically characterized as being prenylated (Glenn et al., 1992). Several others are inferred to be prenylated based on suppression of viral replication by statins (Pronin et al., 2021).

We also evaluated the prenylation status of CaaL/I/M sequences associated with prenylated human proteins. Sequences adhering to consensus motif are considered targets of GGTase-I or interchangeably by FTase or GGTase-I (Krzysiak et al., 2010; Trueblood et al., 1993; Zhang et al., 1994). The CaaX sequences evaluated were from small GTPases (CDC42, RAC1, RAP1B, RHOA and RHOB), G-protein gamma subunits (GBG5 and GBG12), myelin protein CN37, F-box protein FBXL2 and anti-viral protein OAS1 p46 (De Angelis and Braun, 1994; Katayama et al., 1991; Kawata et al., 1990; Kilpatrick and Hildebrandt, 2007; Onono et al., 2010; Soveg et al., 2021; Storck et al., 2019; Wang et al., 2005; Wickenhagen et al., 2021). All Ydj1-CaaL/I/M variants appeared to be fully prenylated in the presence of *Hs*FTase, with exception of CSFL that does not adhere to the consensus sequence due to non-aliphatic amino acids at the a_1_ and a_2_ positions. Of note, a few sequences with non-aliphatic a_1_ amino acids were fully prenylated (i.e., CTII, CKVL, and CQLL). Comparing band mobilities in the presence and absence of *Hs*FTase indicated that several sequences were prenylated in a predominantly *Hs*FTase-dependent manner (CKVL, CLLL, CLVL, CQLL, CSFL, and CVLL). For the remaining sequences, a smaller portion of the population exhibited a mobility shift in the absence of *Hs*FTase (CTII, CIIL, CTIL and CVIL), suggesting that these sequences are either naturally geranylgeranylated naturally farnesylated but strongly subject to alternate prenylation when FTase is compromised (**Figure 4C**).

### *Hs*FTase modifies non-canonical length sequences

The substrate recognition profile of FTase has recently been expanded to certain sequences one amino acid longer or shorter than the tetrapeptide CaaX sequence (i.e., CaX and CaaaX). This expanded specificity was first identified using a combination of yeast genetics and *in vitro* biochemical analyses involving mammalian and yeast FTase (Ashok et al., 2020; Blanden et al., 2018; Schey et al., 2021). Additional CaaaX sequences subject to farnesylation were subsequently identified by *in vitro* screening of synthetic peptide libraries and by searches of the human proteome ((Blanden et al., 2018; Schey et al., 2021) and this study). To extend these observations to *Hs*FTase, previously studied Ydj1-CaX and -CaaaX variants were expressed in the humanized FTase yeast strain and analyzed by gel-shift assay (**Figure 5**). One additional CaX (CHA) sequence was also evaluated due to its association with SARS-CoV2 ORF7b.

**Figure 5.**
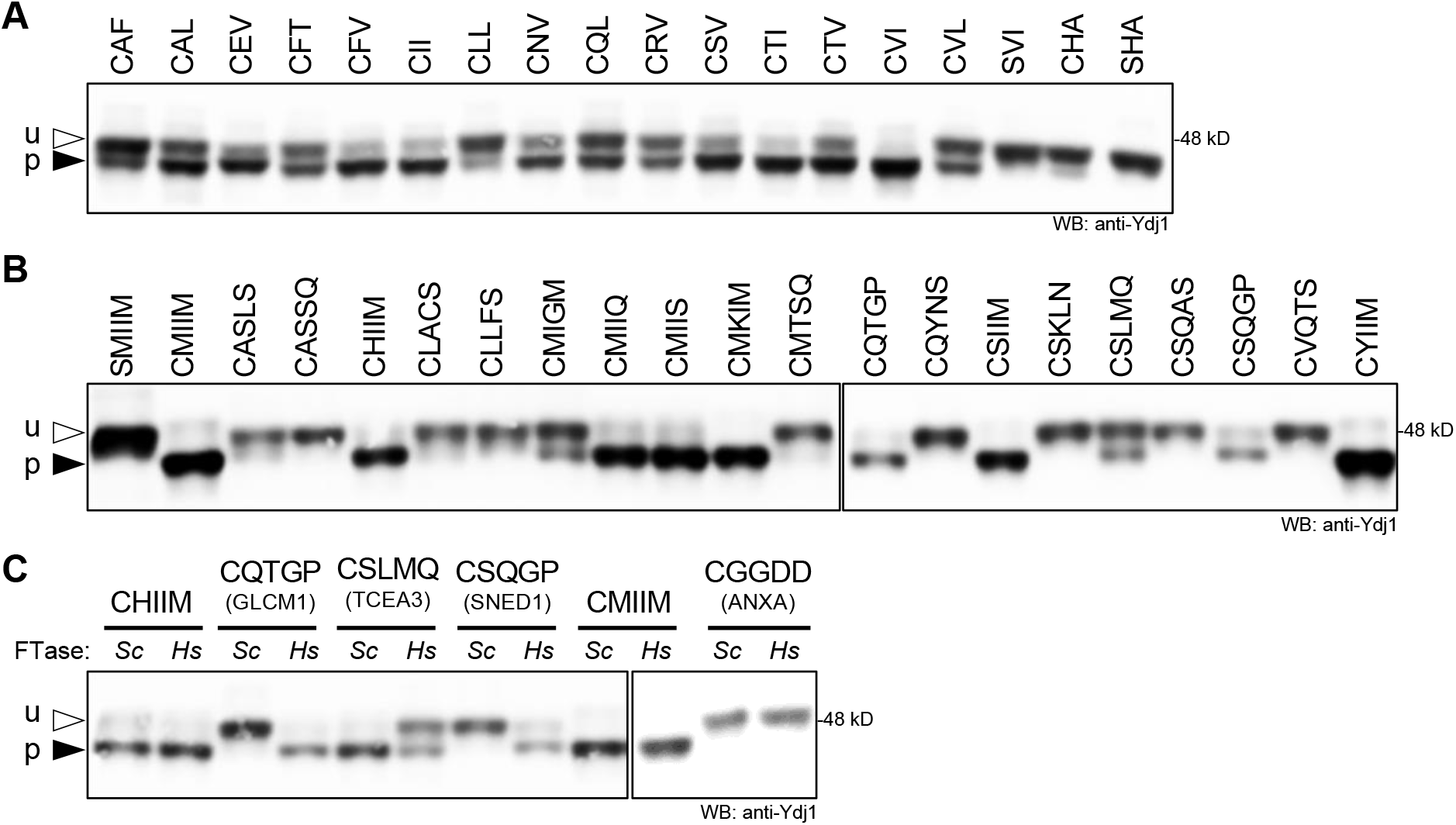
The humanized FTase yeast strain modifies certain non-canonical length sequences. Ydj1-CaaX variants with **A**) shorter CaX and **B**) longer CaaaX sequences were produced in the humanized FTase strain and evaluated by gel-shift assay as described in Figure 3. u – unprenylated Ydj1; p – prenylated Ydj1. **C**) Comparison of Ydj1-CaaaX variant gel mobilities in the presence of *Sc*FTase (*Sc*) and *Hs*FTase (*Hs*). The source of the CaaaX sequence is indicated below each specific sequence. Total cell lysates were prepared and evaluated as described in Figure 3B. Strains used were yWS3186 (*Hs*FTase ydj1Δ) and yWS2544 (*Sc*FTase *ydj1Δ*). Plasmids used are listed in **Supplemental Table S3**. See **Supplemental Table S1** for gel quantification.

The 16 CaX sequences evaluated exhibited varying degrees of farnesylation ranging from >90% (CVI, CTI) to <25% (CLL and CHA), with half exhibiting >50% farnesylation (8 of 16) (**Figure 5A** and **Supplemental Table S1**). The farnesylation patterns were similar to that observed for *Sc*FTase. The very weak modification of Ydj1-CHA, which was quantified to be ∼8% of the population, was not evident with SHA, consistent with the cysteine-dependent nature of this modification, but it is unclear whether such a low level of modification it is likely to be biologically significant for the biology of SARS-CoV2.

The 20 CaaaX sequences evaluated also exhibited varying degrees of farnesylation ranging from >90% (e.g., CMIIM) to unfarnesylated (e.g., CASSQ), with nearly half exhibiting >50% farnesylation (9 of 20) (**Figure 5B**). In most cases, *Sc*FTase and *Hs*FTase reactivities toward Ydj1-CaaX sequences have been highly similar, with the extent of modification being occasionally different for partially modified sequences (**Supplemental Table S1**). Here, however, dramatic differences were observed for a few CaaaX sequences. Two sequences were modified by *Hs*FTase but not *Sc*FTase: CQTGP (associated with GLCM1) and CSQGP (SNED1). One sequence was fully modified by *Sc*FTase but <50% modified by *Hs*FTase: CSLMQ (TCEA3) (**Figure 5C**). The species specificity differences observed with these CaaaX sequences may indicate subtle differences in the FTase substrate binding pockets of these enzymes that warrants future investigation.

This survey of CaX and CaaaX sequences reveals that *Hs*FTase can indeed modify non-canonical length sequences in a cell-based system, albeit only a few exhibit full modification. For the sequences associated with eukaryotic or viral proteins (see **Supplemental Table S5**), it remains to be determined whether any are modified in their native context. Considering that there are 988 CaX and 941 CaaaX human proteins in the UniProtKB/Swiss-Prot database (**Table 1**), it is possible that some of these will be prenylated.

### The humanized FTase yeast strain can be used to identity novel *Hs*FTase target sequences

Two genetic screens, utilizing plasmid libraries based on distinct CaaX protein reporters (i.e., Ras and Ydj1), have comprehensively evaluated CaaX sequence space for reactivity with *Sc*FTase (Kim et al., 2023; Stein et al., 2015). These genetic strategies are fully compatible with the humanized FTase strain developed in this study and could be used to query CaaX sequence space recognized by *Hs*FTase. As proof of principle of this possibility, we performed a miniscreen by transforming the Ydj1-CaaX plasmid library into the humanized FTase strain and recovered a random sampling of thermotolerant and temperature sensitive colonies from which plasmids were recovered, sequenced, and evaluated by thermotolerance and gel-shift assays (**Figure 6** and **Supplemental Table S1**). The results reveal a correlation between the ability to support thermotolerance and being farnesylated by *Hs*FTase. The reciprocate is true for temperature sensitivity and lack of farnesylation, except for the sequences CWIM and CWFC, which were fully farnesylated yet temperature sensitive. The observations for CWIM and CWFC are consistent with the profile of canonical CaaX sequences that temper the ability of Ydj1 to support thermotolerance (i.e., fully-modified; farnesylated, cleaved, and carboxylmethylated) (Berger et al., 2018; Hildebrandt et al., 2016b).

**Figure 6.**
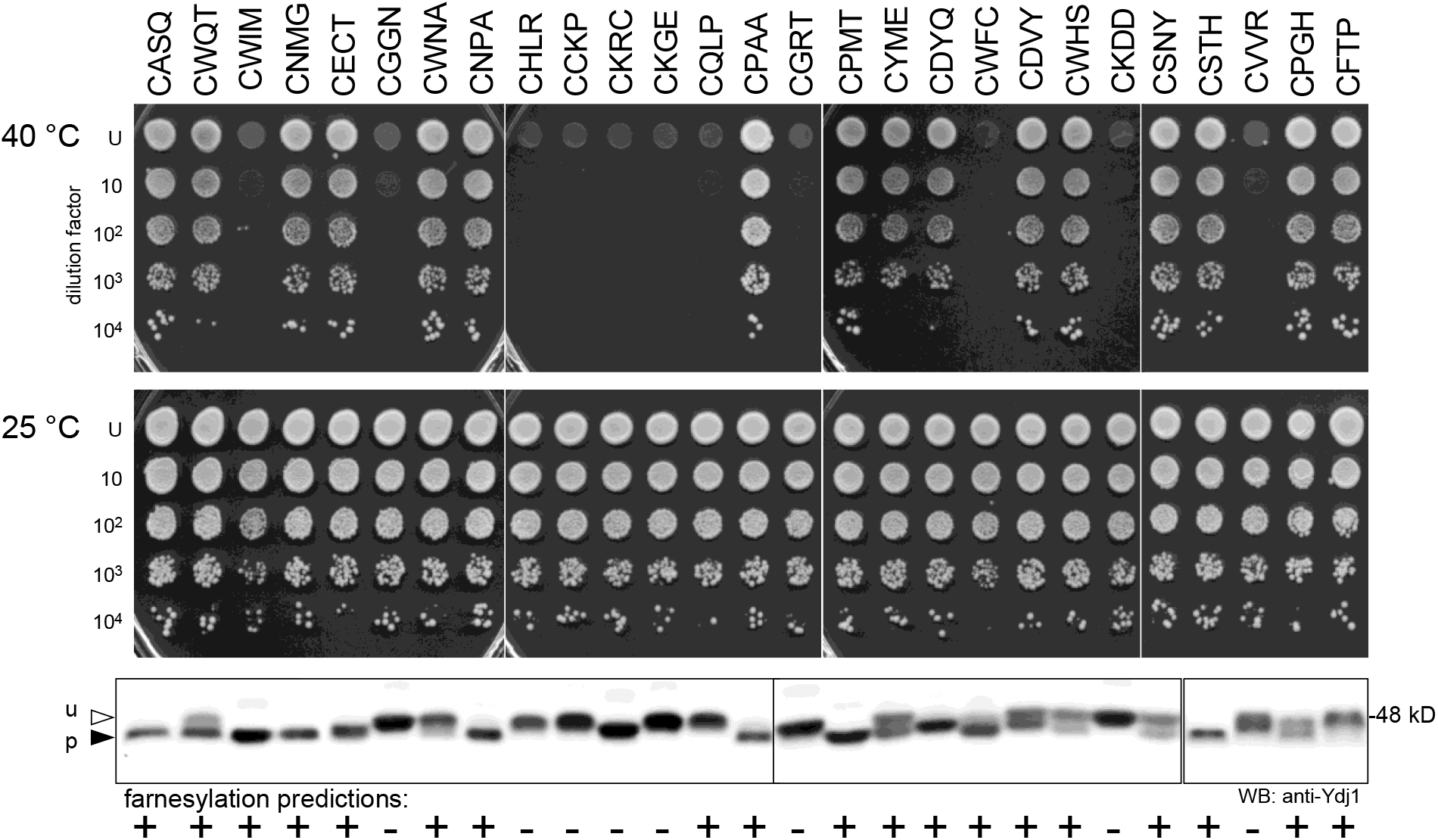
Miniscreen using the humanized FTase strain for identifying prenylatable CaaX sequences. The humanized FTase yeast strain (yWS3186) was transformed with a plasmid library encoding all 8000 Ydj1-CaaX variants. Plasmids were recovered and sequenced from a randomly selected population of thermotolerant and temperature-sensitive colonies. The plasmids were individually retransformed into yWS3186 and resultant strains evaluated by thermotolerance (upper panels), and gel-shift assays (lower panel) as described in Figure 3. Predictions for the farnesylation of each sequence by *Sc*FTase are indicated at the bottom of the figure. Predictions were derived using Heat Map prediction scores reported in Kim et al (Kim et al., 2023); HM score >3 predicts farnesylation, HM score <3 predicts not farnesylated. See **Supplemental Table S1** for gel quantification.

Studies of *Sc*FTase specificity have led to the development of predictive models for prenylation (Berger et al., 2022; Kim et al., 2023). The more recent of these models was used to predict the farnesylation status (positive and negative) of the randomly sampled CaaX sequences. The predictions correlated well with empirical observations for positive and negative modification by *Hs*FTase with the majority of sequences (20 of 26) being accurately predicted using rigorous thresholds (>50% modification as determined by band quantitation was required for positive classification; <10% modification was required for negative classification). For classification purposes, CKRC was judged to be unmodified, consistent with its negative prediction, because the gel-shift associated with this sequence also occurred in the absence of FTase (i.e., *ram1Δ* background), indicating that it is not farnesylated (**Supplemental Figure S5**). Of the outlier sequences, some were predicted to be modified, and indeed exhibited partial modification, but the extent of their modification was below the threshold for positive classification (i.e., CWNA, CWHS, CSNY and CPGH). By contrast, two outlier sequences that were predicted to be modified were unmodified (i.e., CQLP and CFTP), perhaps indicating that a terminal proline is unfavorable for farnesylation by *Hs*FTase, which has been previously reported (Kim et al., 2023; Moores et al., 1991).

Among the randomly sampled sequences, only CPAA was found to exist in the human proteome when querying the UniprotKB/SwissProt database. It is associated with two proteins: the ER-localized DNase-1-like protein (P49184) and the extracellular laminin alpha subunit (Q16363). While CPAA was reactive with *Hs*FTase, the subcellular locations of these human proteins likely disqualify them from being substrates of cytosolic FTase. Nevertheless, our results suggest ∼77% accuracy when using a farnesylation prediction model developed using *Sc*FTase specificity data to predict modification by *Hs*FTase (Kim et al., 2023). The substrate specificities of *Sc*FTase and *Hs*FTase, although highly similar, are not identical, which prompts the need for future studies aimed at fully evaluating CaaX sequence space in the context of *Hs*FTase. Such studies are now possible with our humanized FTase yeast strain, and such investigations may better refine the target profile of *Hs*FTase, potentially leading to the discovery of novel CaaX proteins.

### Interspecies complementation analysis of GGTase-I subunits

To develop a yeast system for expressing *Hs*GGTase-I, studies were first performed to assess the functional equivalence of *Sc*GGTase-I and *Hs*GGTase-I subunits. Because GGTase-I activity is essential for yeast viability, strains lacking a *Sc*GGTase-I subunit can only be propagated when complemented by plasmid-encoded copy of the missing gene (e.g., *ram2*Δ [*URA3 RAM2*]).

These strain backgrounds were further modified to introduce plasmid-encoded copies of one or both orthologous *Hs*GGTase-I subunit genes FNTA and PGGT1B under different promoter conditions. The strains were cultured, serially diluted, and spotted onto media containing 5-fluroorotic acid (5FOA) that counter selects for the *URA3*-marked plasmids encoding *RAM2* or *CDC43* in the parent strains. This strategy forced strains to contain either species matched or mismatched subunits of GGTase-I in the absence or presence of FTase (*ram2*Δ and *cdc43*Δ backgrounds, respectively).

The 5FOA analysis revealed, as observed for FTase, that neither FNTA nor PGGT1B alone could complement for the absence of its yeast ortholog *ram2Δ* or *cdc43*Δ, respectively, indicating that interspecies subunits do not form functional GGTase-I complexes (**Figure 7A**, 5FOA, 25 °C). This was observed whether FNTA and CDC43 (conditions **4** and **5**) or Ram2 and PGGT1B (**8** and **9**) were co-expressed, and the result did not change when human subunits were over-expressed using the *PGK1* strong constitutive promoter (**5** and **8**). Only when strains producing both *Hs*GGTase-I subunits, whether expressed from the yeast orthologous promoters (**3** and **7**) or the *PGK1* promoter (**2** and **6**), was growth equivalent to wildtype yeast (**1**). This observation contrasts with the FTase situation where both *Hs*FTase subunits had to be expressed from the *PGK1* promoter for functional complementation (**Supplemental Figure S1**). As additional confirmation that yeast growth was specifically due to *Hs*GGTase-I activity, we observed that *Hs*GGTase-I supported growth on 5FOA at 37 °C in the *cdc43*Δ but not *ram2Δ* background (**Figure 7A**, 37 °C). FTase activity is required for growth at higher temperature (He et al., 1991). The *cdc43*Δ strains (**6** and **7**) grew because they retain endogenous FTase activity (i.e., Ram2/Ram1 complex), while the *ram2*Δ strains (**2** and **3**) failed to grow because they lack FTase activity (i.e., Ram1 subunit only). Thus, it can be inferred that growth on 5FOA at 25 °C in the *ram2*Δ background can be attributed entirely to *Hs*GGTase-I.

**Figure 7.**
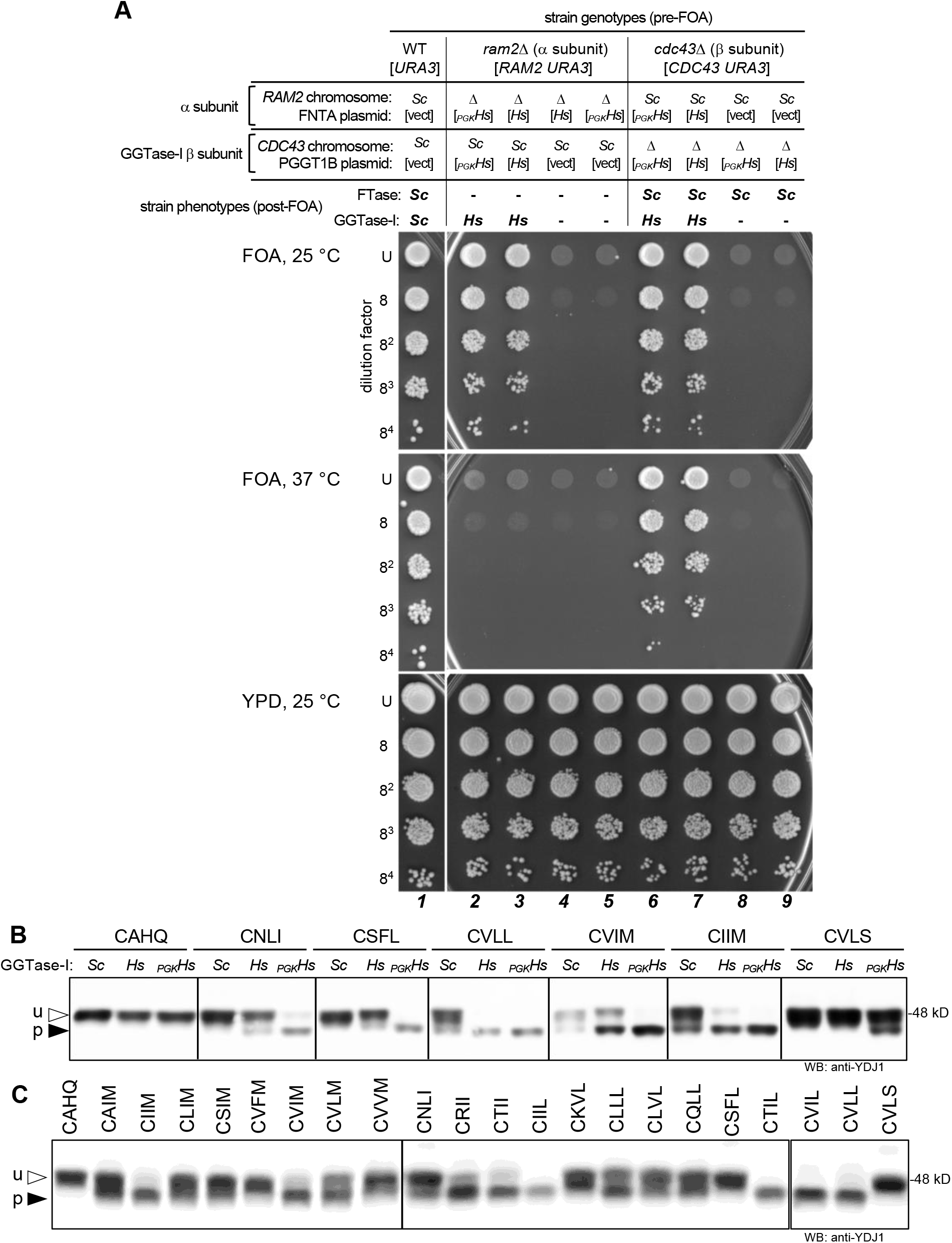
Interspecies complementation studies of GGTase-I subunits. *Hs*GGTase-I and *Sc*GGTase-I subunits are not interchangeable, but co-expression of both α and β subunits of *Hs*GGTase-I in yeast can restore GGTase-I activity in the absence of *Sc*GGTase-I. **A**) Viability assay to assess GGTase-I-dependent growth. The yeast genetic backgrounds have only one of the naturally encoded *Sc*GGTase-I subunits encoded on the chromosome, with the presence (*Sc*) and absence (Δ) of the specific subunit indicated above the panel. Because GGTase-I activity is essential for yeast viability, the other subunit is plasmid-encoded. The strains also contain plasmid(s) encoding *Hs*GGTase-I FNTA and/or PGGT1B subunits (indicated by brackets) driven by either the orthologous yeast gene promoter (*Hs*) or the *PGK1* promoter (*_PGK_Hs)*. Select strains were also transformed with empty vectors ([vect]) so that all strains had the same selectable markers. The pre-FOA strain genotypes are indicated at the top of the panel. Strains were cultured to the same density, serially diluted, and the dilution series mixtures spotted onto YPD and 5FOA media. Growth on YPD media provides an assessment for the quality of the serial dilutions. 5FOA media counter selects for the *URA3* gene and effectively eliminates the *URA3*-marked plasmid, resulting in the post-FOA phenotypes indicated at the top of the panel. Growth on 5FOA at 25 °C indicates the presence of functional GGTase-I (**2**, **3**, **6** and **7**). Growth on 5FOA at 37 °C is also dependent on functional FTase, which requires α and FTase β subunits of the same species (**6** and **7**). Strains used were yWS3481 (***1***), yWS3388 (***2***), yWS3414 (***3***), yWS3639 (***4***), yWS3287 (***5***), yWS3387 (***6***), yWS3413 (***7***), yWS3285 (***8***), and yWS3638 (***9***). **B)** Gel-shift analysis of Ydj1-CaaX variants produced in yeast expressing *Sc*GGTase-I (*Sc*) or plasmid-encoded *Hs*GGTase-I (*Hs* or *_PGK_Hs*). Total cell lysates were prepared and analyzed as described in Figure 3B. Strains used were yWS3209 (*Sc*), yWS3451 (*Hs*), and yWS3169 (*_PGK1_Hs*). **C)** The humanized GGTase-I strain allows prenylation of Ydj1-CaaM/I/L variants taken from human proteins. The expression of GGTase-I subunits in this strain was driven from the orthologous yeast gene promoters to better match the natural *in vivo* activity of yeast GGTase-I. Total cell lysates were prepared and analyzed as described in Figure 3B. The strain used was yWS3451. The plasmids in used in this figure are listed in **Supplemental Table S3**. See **Supplemental Table S1** for gel quantification.

### *Hs*GGTase-I expressed in yeast modifies both canonical and non-canonical CaaX sequences

Evidence suggests that GGTase-I may have broader substrate specificity at the a_1_ and a_2_ positions akin to what has been observed for yeast and mammalian FTase (Kawata et al., 1990; Kilpatrick and Hildebrandt, 2007; Lebowitz et al., 1997; Onono et al., 2010). We investigated this issue more thoroughly using a three-plasmid system to examine geranylgeranylation of plasmid-encoded Ydj1-CaaX variants in the context of *Hs*GGTase-I (**Figure 7B**). The strains evaluated all lack endogenous FTase and Ydj1, but express either endogenous GGTase-I (*Sc*) or plasmid-encoded *Hs*GGTase-I subunits from the orthologous *RAM2* and *CDC43* promoters (*Hs*) or constitutive *PGK1* promoters (*_PGK_Hs*). Gel-shift analysis revealed that neither *Sc*GGTase-I nor *Hs*GGTase-I were able to modify Ydj1 harboring the CAHQ sequence that is associated with the Ydj1 human ortholog DNAJA2 that is strictly a farnesylated sequence (Andres et al., 1997). Other CaaX sequences (CNLI, CSFL, CVLL, CVIM, CIIM and CVLS) exhibited no or limited modification by *Sc*GGTase-I, yet were better modified by *Hs*GGTase-I. This was more evident when *Hs*GGTase-I subunit pairs were expressed using the strong *PGK1* promoter. These observations suggest that *Hs*GGTase-I expressed in this yeast system is either more active or has a different specificity than *Sc*GGTase-I, and that *Hs*GGTase-I indeed has the potential to modify certain non-canonical CaaL/I/M sequences (e.g., CNLI and CSFL).

To further survey the geranylgeranylation potential of CaaX sequences, we evaluated a test set of sequences that included 22 sequences that mostly matched the CaaL/I/M consensus. For this analysis, we used the yeast system producing *Hs*GGTase-I from orthologous yeast promoters to reduce the potential of over-expression artifacts (**Supplemental Figure S2C**). Most sequences exhibited a gel-shift indicative of modification, but only a few exhibited complete modification (i.e., CIIL, CTIL, CVIL, CVLL) or near-complete modification (CIIM, CVIM, CTII) (**Figure 7C**). Within this data set, we observed that the a_1_ position influenced prenylation by *Hs*GGTase-I. This is evident when comparing the CxIM set of sequences: CSIM, CAIM, and CLIM (∼50% modified) vs. CIIM and CVIM (100% modified). A similar observation was made when comparing CxLL sequences: CLLL and CQLL (∼50% modified) vs. CVLL (100% modified). Sequences where the X position did not match the CaaL/I/M consensus (CAHQ, CVLS) were unmodified by *Hs*GGTase-I.

### Human lamins and Pex19 CaaX sequences have distinct CaaX protease profiles

The ability to humanize yeast for investigations of CaaX protein modifications can be extended to studies of the human CaaX protease Rce1. This protease cleaves CaaX proteins that follow the canonical modification pathway, which are subject to the coupled modifications of proteolysis and carboxymethylation. Rce1 prefers to cleave prenylated CaaX sequences having an aliphatic amino acid at a_2_, and this substrate specificity is generally conserved among orthologs (Mokry et al., 2009; Plummer et al., 2006). The Ste24 protease has also been referred to as a CaaX protease, but its primary role is in protein quality control, and its substrates do not need to be prenylated, unlike Rce1 (Ast et al., 2016; Boyartchuk et al., 1997; Hildebrandt et al., 2016a; Runnebohm et al., 2020).

In yeast, the **a**-factor mating pheromone is a useful reporter of CaaX proteolysis that can be coupled with expression of human Rce1 and ZMPSte24 for studies of target specificity in a cell-based model (Mokry et al., 2009; Plummer et al., 2006). Such studies can be used to help resolve the CaaX protease preferences of medically relevant human proteins. For example, defects in the proteolytic processing of Prelamin A (CSIM) are associated with progeria-like diseases (Barrowman et al., 2012b). Prelamin A is subject to two proteolytic events that yield Lamin A: one within the CaaX sequence and another 15 amino acids from the C-terminus. Both cleavage events have been attributed to ZMPSte24 (Barrowman et al., 2012a; Quigley et al., 2013). With a humanized CaaX protease yeast model, however, we observe that **a**-factor-CSIM is cleaved by *Hs*Rce1, not ZMPSte24, in accordance with more recent observations (**Figure 8**) (Berger et al., 2022; Nie et al., 2020). We extended these studies to evaluate the cleavage of the Lamin B CaaX sequence (CAIM), which was cleaved by both *Hs*Rce1 and ZMPSte24. This result suggests that FTase inhibitors (FTIs) may impact the CaaX processing of both lamins, while Rce1 inhibitors are likely to differentially impact Prelamin A and Lamin B processing. We also examined cleavage of the human peroxisomal chaperone Pex19 CaaX sequence. Farnesylation of Pex19 is critical for its interaction with client proteins, and mutations of Pex19 unrelated to its CaaX sequence are associated with Zellweger syndrome (Emmanouilidis et al., 2017; Matsuzono et al., 1999). Human Pex19 (CLIM) and yeast Pex19 (CKQQ) have strikingly different CaaX sequences despite moderate homology (i.e., 21% identity; 48% similarity) (Madeira et al., 2022). Both sequences are fully prenylated in the context of either *Hs*FTase or *Sc*FTase (**Figure 4A** and **Supplemental Figure S4**). Using **a**-factor-CaaX variants harboring these sequences, CLIM was observed to be cleaved by both *Hs*RCE1 and ZMPSte24, whereas the non-canonical CKQQ sequence was not. This result suggests that Pex19 proteins have evolved to have C-termini with distinct biophysical properties: canonical for human Pex19 and only isoprenylated for yeast Pex19. The underlying reasons for these differences remain unknown. Nonetheless, our studies with lamins and Pex19 fully support the utility of the humanized CaaX protease model system for expanding studies of CaaX protein post-translational modifications (PTMs) beyond the initial prenylation step. Such studies could prove useful for predicting the potential effects of inhibiting the Rce1 CaaX protease, which would impact a narrower range of CaaX protein targets than prenyltransferase inhibitors.

**Figure 8.**
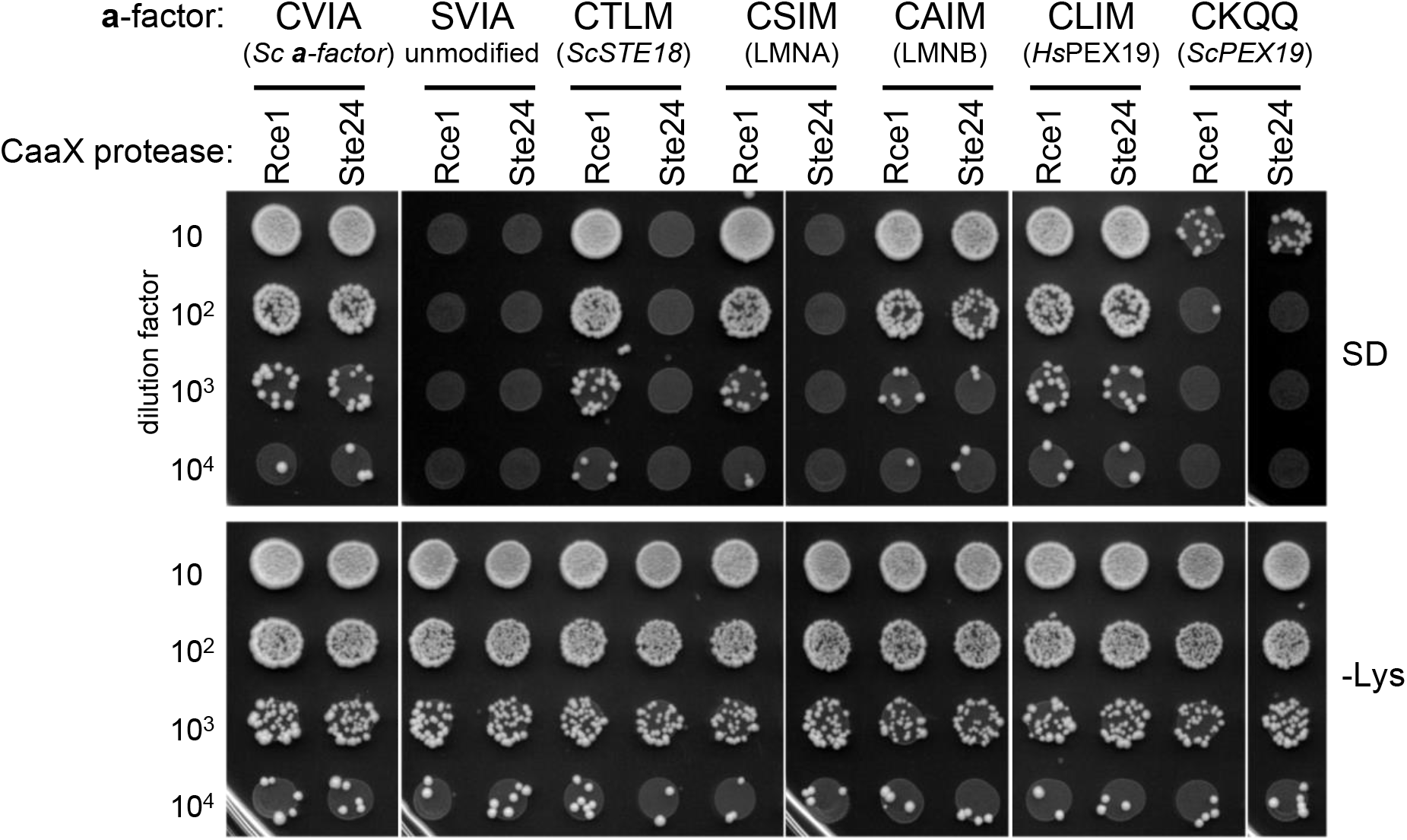
Prenylation studies can be coupled with CaaX proteolysis studies to better understand CaaX PTMs. The mating assay was performed as described in Figure 1B using *MAT***a** strains that carry plasmid-encoded **a**-factor-CaaX variants and plasmid-encoded *Hs*RCE1 (Rce1) or ZMPSTE24 (Ste24) CaaX proteases. The source of the CaaX sequence is indicated below each specific sequence. The strain background for this experiment lacks endogenous **a**-factor (*MFA1* and *MFA2*) and yeast CaaX protease genes (yWS164; *MAT***a** *mfa1Δ mfa2Δ rce1Δ ste24Δ*). *MAT***a** strains were mixed with *MAT*α cells (SM1068), mixes subject to 10-fold dilutions, and dilution mixtures spotted onto minimal media (SD) and SC-lysine (-Lys) media. Growth on SD indicates diploid formation, which is a direct indicator of **a**-factor mating pheromone production. Growth on –Lys reflects the input of *MAT***a** cells. Plasmids are listed in **Supplemental Table S3**.

## Discussion

We report the development of yeast strains that can be used for systematic characterization of prenylation by human FTase and GGTase-I. The humanized FTase strain, with subunits FNTA and FNTB integrated into the yeast genome, was phenotypically equivalent to yeast expressing native FTase in four functional tests: 1) production of **a**-factor mating pheromone, 2) Ras2 localization 3) Ydj1-based thermotolerance, and 4) Ydj1 farnesylation. Thus, the humanized FTase yeast strain provides *in vivo* FTase activity levels compatible for relatively rapid characterization of candidate CaaX sequences and heterologously expressed human CaaX proteins that may be targets for farnesylation. Moreover, the humanized FTase yeast strain provides a cell-based system that overcomes concerns associated with *in vitro* farnesylation assays that depend on chemically modified substrates and unnatural enzyme and substrate amounts. For example, the annexin A2 CGGDD extended CaaaX sequence has been reported as farnesylated with methods involving metabolic labeling with an azido-farnesyl analog (Kho et al., 2004), but this sequence is unmodified by human FTase in our cell-based system (**Figure 5C**). The convenience of the yeast system for heterologous protein expression makes it ideal for coupling with other methods, such as mass-spectrometry, for confidently validating the prenylation status of candidate CaaX proteins.

Limited comparative data exists for the specificities of mammalian FTase relative to *Sc*FTase. *In vitro,* rat FTase and *Sc*FTase and exhibit similar specificities when evaluated against a CVa_2_X peptide library (Wang et al., 2014). Our *in vivo* results confirm and further extend such observations to reveal that *Hs*FTase expressed in yeast and native *Sc*FTase have a striking degree of substrate conservation (see **Supplemental Figure S4**, **Supplemental Table S1**). This is most evident in the ability of both enzymes to similarly prenylate canonical and prenylation-only CaaX sequences. Thus, *Hs*FTase also appears to possess the broadened ability to modify non-canonical sequences (i.e., sequences without aliphatic amino acids at the a_1_ and a_2_ positions), which was initially reported for *Sc*FTase (Berger et al., 2018). These findings indicate that farnesylation prediction algorithms developed using *Sc*FTase should work reasonably well for predicting *Hs*FTase specificity (Kim et al., 2023). The conservation between human and yeast FTases extends to prenylation of non-standard length sequences that are one amino acid shorter or longer than the conventional tetrapeptide CaaX sequence, although species specific differences in substrate recognition of longer CaaaX sequences was observed.

The humanized GGTase-I strain, with subunits FNTA and PGGT1B expressed from low copy plasmids and driven by the orthologous yeast promoters, was phenotypically equivalent to yeast expressing native GGTase-I in supporting viability. The humanized GGTase-I strain, however, displayed more robust activity than endogenous yeast GGTase-I in modifying Ydj1-CaaL/I/M variants. This observation could be due to higher GGTase-I enzyme levels and/or subtle specificity differences between yeast and human enzymes. Human GGTase-I prenylated a wide range of CaaL/I/M sequences and did not prenylate CaaS/Q sequences, in alignment with expectations. One feature that was conserved across species was the ability of GGTase-I to alternatively prenylate normally farnesylated sequences. This typically occurs when FTase activity is reduced through action of FTIs, and in our case, when the FTase activity was genetically ablated. The observation that yeast and human FTase and GGTase-I can alternatively prenylate Ras sequences CIIM and CVIM when using Ydj1 or GFP-Ras2 reporters (**Supplemental Figure S6**) demonstrates that recognition of these sequences by both prenyltransferases is transferable and evolutionarily conserved between yeast and human enzymes, and likely across other species as well.

The scope of potentially prenylated proteins in protein databases is expansive with over 6000 annotated proteins in eukaryotes and viruses ending in Cxxx. When expanded to 3-mer and 5-mer length sequences, this number is over 15,000 (see **Table 1**). Not accounted for in this table, is the potential number of prokaryotic CaaX proteins. While prokaryotes lack protein prenylation machinery, there is evidence that some pathogenic bacteria, such as *Legionella pneumophila* and *Salmonella typhimurium,* encode CaaX proteins that are prenylated by host enzymes as part of their infectious life cycle (Ivanov et al., 2010; Reinicke et al., 2005). Thus, a reliable prediction algorithm and reporter systems to identify and verify the subpopulation of CaaX sequences that are likely to be modified by either FTase or GGTase-I will be a key step toward defining the prenylome across many different organisms.

Viral genomes encode many CaaX proteins that might be prenylated by human FTase and/or GGTase-I. We have evaluated several such sequences using our system, with results confirming instances of previously reported prenylation or indicating compatibility for prenylation. For example, we observed that the CaaX sequence of Hepatitis Delta Virus Large Delta antigen (L-HDAg; CRPQ) is modified by human and yeast FTases, but not by GGTase-I. Moreover, we observed that the CRPQ sequence is neither cleaved nor carboxylmethylated, consistent with its non-canonical sequence (see **Figure 4** and **Supplemental Figures 2 and 3**). Prenylation of L-HDAg is required for virion assembly, and the FTI Lonafarnib is currently in Phase 3 clinical trials for use in combination treatments for Hepatitis D infections (Khalfi et al., 2023; Koh et al., 2015; Yardeni et al., 2022). We also determined that the CaaX sequence of human cytomegalovirus protein of unknown function IRL9 (CRIQ) can be farnesylated, cleaved and carboxymethylated. The modifications of IRL9 have not been previously reported, so it is a good candidate for additional investigations related to its prenylation status and impact of PTMs to virion formation. Other viral CaaX sequences CRII (Influenza virus H1N1 protein PB2), CVLM (Molluscum contagiosum virus protein MC155R) and CNLI (ASFV protein A179L) were substrates of both *Hs*FTase and *Hs*GGTase-I in our system. If these sequences are prenylated in their native context, dual prenyltransferase inhibitors might have better utility for blocking modification. Given that all the viral CaaX sequences evaluated in our study were targets for prenylation, it is prudent to consider the role that the prenylation of viral proteins may have on viral propagation and infection. The ability to readily assess the prenylation status of viral CaaX sequences afforded by our cell-based system could provide the necessary evidence for new investigations of viral biology or viral therapies in human and veterinary medicine involving FTIs, geranylgeranyltransferase inhibitors (GGTIs), dual prenyltransferase inhibitors (DPIs), and possibly CaaX protease and ICMT inhibitors as well.

Our system is well-suited for assessing the prenylation status of human proteins as supported by the modification of heterologously expressed human HRas61 and DNAJA2. An important caveat to consider is that SDS-PAGE gel-shifts do not universally occur for all prenylated proteins, which is why Ydj1 is such a great reporter protein for testing CaaX sequences. Unfortunately, our system does not exclude the possibility of a CaaX sequence being reactive in the context of Ydj1, but unreactive its native context, for a myriad of reasons (e.g., structurally inaccessible C-terminus, localization of protein inaccessible to cytosolic prenyltransferases, alternate cysteine modifications such as disulfide bonding, etc.). Therefore, positive results with our system should be coupled with additional validation of prenylation in native contexts. Nonetheless, the ease and convenience of the humanized FTase and GGTase-I yeast strains that we have developed are an important and valuable new resource for initial investigations into evaluating the prenylation potential of a wide array of CaaX sequences.

## Materials and Methods

### Yeast strains

The humanized prenyltransferase strains are summarized in **Supplemental Figure S2**. All yeast strains used in this study are listed in **Supplemental Table S2**. Detailed descriptions of strain constructions can be found in the **Supplemental Methods** file. Yeast were typically cultured in standard liquid and solid yeast media unless otherwise noted. For expression of Myc-HRas61 from the inducible *MET25* promoter (p-05547) in methionine auxotroph strains (i.e., yWS2544 and yWS3186), yeast were first cultured in SC-leucine containing 20 µg/ml methionine to late log phase, washed, and resuspended in SC-leucine containing 2 µg/ml methionine to allow for both growth and induction of the *MET25* promoter.

New strains were typically created for this study by standard genetic manipulations starting with commercially available haploid and heterozygous diploid genomic deletions. *KAN^R^* and *NAT^R^* marked gene replacements were confirmed by growth on YPD containing G418 (200 µg/ml; Research Products International) or nourseothricin (100 µg/ml; GoldBio), respectively. All gene replacements were further checked by PCR to confirm the presence of the knockout at the correct locus and absence of the wild-type open reading frame. Plasmids were introduced into strains via a lithium acetate-based transformation procedure (Elble, 1992). Yeast sporulation was carried out in a solution of 2% potassium acetate, 0.25% yeast extract, and 0.1% dextrose. Spore enrichment and random spore analysis followed published methods (Rockmill et al., 1991). For the humanized FTase strains (yWS3186 and yWS3220), *P_PGK1_-FNTB* was integrated at the *RAM1* locus, replacing the open reading frame, using a loop-in loop out strategy, and *P_PGK1_-FNTA* was integrated at the *his3Δ1* locus by homologous recombination using a *HIS3*-based integrative plasmid (Schey et al., 2023).

### Plasmids

The plasmids used in this study are listed in **Supplemental Table S3**. Detailed descriptions of plasmid construction can be found in the Supplemental Methods file. Plasmids were analyzed by diagnostic restriction digest and DNA sequencing (Eurofins Genomics, Louisville, KY) to verify the entire open reading frame and surrounding sequence. Plasmids recovered from the Ydj1-CaaX Trimer20 library are described elsewhere (Kim et al., 2023). New plasmids encoding Ydj1-CaaX and **a**-factor-CaaX (encoded by *MFA1* gene) variants were made by PCR-directed plasmid based recombinational cloning as previously described (Berger et al., 2018; Oldenburg et al., 1997). Plasmids with GFP-Ras2-CaaX variants were generated by QuikChange site-directed mutagenesis. Plasmids encoding *Hs*FTase and *Hs*GGTase-I subunits were created in multiple steps. First, the open reading frames (ORFs) and flanking 5′ and 3′ sequences of yeast prenyltransferase subunits were PCR-amplified from strain BY4741 and subcloned into the multicloning sites of appropriate pRS series vectors (Sikorski and Hieter, 1989). In parallel, synthetic cDNAs for human FNTA, FNTB and PGGT1B that were codon optimized for *S. cerevisiae* were subcloned into the PstI and XhoI sites of pBluescriptII KS(-); all cDNAs were commercially obtained (GenScript). The human ORFs were engineered to also have 39 bp of flanking sequence on both the 5′ and 3′ends that equivalently matched the flanking sequences of the orthologous yeast ORFs. Next, DNA fragments encoding the human ORFs and yeast flanking sequences were used with recombination-based methods for direct gene replacement of the plasmid-encoded yeast ORFs. The *PGK1* promoter was amplified and used to replace the orthologous yeast promoters by similar recombination-based methods as described in **Supplemental Materials**. To change selectable markers, the various FTase and GGTase-I promoter-containing segments were subcloned as XhoI-SacI fragments into different pRS vector backbones by conventional ligation-based cloning.

### Temperature sensitivity assay

Thermotolerance assays were performed as previously described (Blanden et al., 2018; Hildebrandt et al., 2016b). In brief, plasmid-transformed strains were cultured in appropriate selective synthetic drop-out (SC-) liquid media to saturation (25 °C, 24 hours); strains without plasmids were cultured in SC complete media. Cultures were serially diluted into H_2_O (10-fold dilutions), and dilutions replica pinned onto YPD plates. Plates were incubated (25 °C for 72 hours; 37 °C for 48 hours; 40 °C and 41 °C for 72 hours) then digitally imaged using a Canon flat-bed scanner (300 dpi; grayscale; TIFF format).

### Yeast lysate preparations and immunoblots

Cell-mass equivalents of log-phase yeast (A_600_ 0.95-1.1) cultured in appropriate selective SC-liquid media were harvested by centrifugation, washed with water, and processed by alkaline hydrolysis and TCA precipitation (Kim et al., 2005). Total protein precipitates were resuspended in urea-containing Sample Buffer (250 mM Tris, 6 M urea, 5% β-mercaptoethanol, 4% SDS, 0.01% bromophenol blue, pH 8), and analyzed by SDS-PAGE (9.5%) followed by immunoblotting. Blots were processed according to standard protocols. Antibodies used for Western blots were rabbit anti-Ydj1 polyclonal (1:10,000 dilution; gift from A. Caplan); mouse anti-c-Myc 9E10 monoclonal (1:1000 dilution, Santa Cruz Biotech. #sc-40); mouse anti-GFP B-2 monoclonal (1:500, Santa Cruz Biotech. # sc-9996); mouse anti-DnaJA2(7) monoclonal (1:500 dilution, Santa Cruz Biotech. sc-136515); mouse anti-HIS his.h8 monoclonal (1:1000 dilution, VWR #101981-852); goat HRP-anti-rabbit and goat HRP-anti-mouse (1:1000 dilutions, Kindle Biosciences, LLC #R1006 and R1005, respectively). Immunoblots were developed with ECL reagent (ProSignal Pico ECL Spray or Kindle Biosciences KwikQuant Western Blot Detection Kit) and images were digitally captured using the KwikQuant Imager system (Kindle Biosciences). Adobe Photoshop was used for cropping and rotating of images; no other image adjustments were applied.

Prenylation of Ydj1-CaaX was quantified using ImageJ and multiple exposures having good dynamic range of band intensities. Percent prenylation was calculated as the intensity of the lower (prenylated band) divided by total intensity of the prenylated and unprenylated (upper) bands. Data presented in **Supplemental Table S1** represent the averages of biological replicates. For duplicate samples, values are reported as an average and associated range. For multiplicate samples, the values are reported as an average and standard error of the mean (SEM).

### Yeast mating assay

Qualitative and quantitative yeast mating assays were performed as detailed previously (Berger et al., 2018). In brief, *MAT***a** test strains and the *MAT*α strain (IH1793) were cultured to saturation at 25 °C in appropriate selective SC-media and YPD liquid media, respectively. Cultures were normalized by dilution with fresh media (A_600_ 1.0), mixed 1:9 (*MAT***a**:*MAT*α) in individual wells of a 96-well plate, and cell mixtures subject to 10-fold serial dilution using the *MAT*α cell suspension as the diluent. For qualitative analyses, each dilution series was replica pinned onto SC-lysine and minimal SD solid media. For quantitative analysis, a portion of a dilution mixture was spread onto SC-lysine and SD plates in duplicate, and colonies counted after plate incubation (72 hours, 30 °C). The SC-lysine cell count reports on the total number of *MAT***a** haploid cells initially in the mixture while the SD cell count reports on the number of mating events (i.e., diploids). Mating efficiencies were normalized to an **a**-factor (*MFA1)* positive control within each experiment.

### Microscopy

Imaging was performed as detailed previously (Ravishankar et al., 2023). In brief, yeast transformed with *CEN URA3 P_Ydj1_-GFP-RAS2-CaaX* variants were cultured to late log (A_600_ 0.8-1) in SC-uracil liquid media and viewed using a Zeiss Axio Observer microscope equipped with fluorescence optics (Plan Apochromat 63X/1.4 N.A objective). Images were captured using AxioVision software and minor image adjustments performed using Adobe Photoshop.

### Humanized yeast miniscreen for identification of Ydj1-CaaX variants

yWS3186 was transformed with the Ydj1-CaaX plasmid library (pWS1775), plated onto SC-uracil solid media, and incubated 4 days at 25 °C. Single colonies of varying size were individually cultured and subject to the temperature sensitivity assay described above except that 3 spots of 20-fold serial dilutions were prepared. Growth at 41 °C was scored on a scale of 0 to 5 relative to control strains yWS3311 (Ydj1-CASQ, score of 5) and yWS3312 (Ydj1-SASQ, score of 0). Plasmids were recovered from candidates scoring 0 (n=11) and candidates scoring 5 (n=15), sequenced to identify the CaaX sequence, and retransformed into yWS3186 for retesting by the temperature sensitivity assay (10-fold dilutions) and SDS-PAGE gel-shift assay.

### 5-fluroorotic acid (5FOA) assay

BY4741, yWS3106 and yWS3109 were transformed with combinations of *HIS3* and *LEU2* based plasmids carrying *Hs*FNTA, *HsP*GGT1B, and/or empty vector plasmids such that all strains had the same selectable markers. Strains cultured on SC-histidine-leucine-uracil solid media were used to inoculate SC-histidine-leucine liquid media. After incubation (25 °C, 24 hours), cultures were normalized with fresh media (A_600_ 2.0), added to wells of a 96 well plate, serial diluted 8-fold with H_2_O as the diluent, and the dilution series pinned onto SC complete solid media containing 1 mg/ml 5-fluroorotic acid (5FOA).

### Search for CaaX sequences

Scan Prosite (https://prosite.expasy.org/scanprosite/) was used to search for CaaX sequences in *Homo sapiens* and viruses in the UniProtKB/Swiss-Prot database using the search strings “CXXX>”, “C{C}XXX>”, or “{C}CXX>” (search performed 07/12/2023). For certain CaaaX and CaX and sequences that were studied in previous publications but have no human or virus associated protein in the annotated UniProtKB/Swiss-Prot database, the search was expanded UniProtKB/TrEMBL database, inclusive of all eukaryotes.

## Footnotes

1 the term CaaX will refer to all C-terminal tetrapeptide sequences whether or not they fit the traditional consensus sequence.

## Acknowledgements

We thank Dr. Avrom Caplan (City College of New York) for anti-Ydj1 antibody, Dr. Jun Wang (Institute Pasteur of Shanghai, Chinese Academy of Sciences) for discussions on ASFV CaaX proteins, and Schmidt lab members for constructive feedback and technical assistance.

## Data Availability

Strains and plasmids generated for this study are available upon request.

